# Increased proportion of growth-arrested bacilli in acidic pH adaptation promotes *Mycobacterium tuberculosis* treatment survival

**DOI:** 10.1101/2025.07.14.664820

**Authors:** Eun Seon Chung, William C. Johnson, Maliwan Kamkaew, Morgan E. McNellis, Trever C. Smith, Srinivasan Vijay, Nguyen Thuy Thuong Thuong, Shumin Tan, Bree B. Aldridge

## Abstract

The ability of *Mycobacterium tuberculosis* (Mtb) to adapt its growth behavior in response to host environments promotes survival against immune and drug stressors. However, how these behaviors shift at the single-cell level remains poorly understood. Here, we show that Mtb adapts to acidic conditions by increasing the proportion of bacteria in a growth-arrested state, rather than uniformly slowing the growth rate of the entire population. This non-growing subpopulation exhibits greater tolerance to ethambutol. Clinical strains displayed higher proportions of growth-arrested cells under both neutral and acidic conditions, suggesting that growth arrest serves as a bet-hedging strategy during infection. Though the PhoPR two-component system contributes to regulating this non-growing state, we show that it is a partial regulator of the non-growing bacterial subpopulation and that additional transcriptional regulators are involved. Our study demonstrates that non-growing subpopulations of Mtb provide fitness benefits and are an active adaptation to environmental cues and not a passive consequence of stressors.

## INTRODUCTION

Tuberculosis (TB), caused by infection with the bacterium *Mycobacterium tuberculosis* (Mtb), requires a lengthy multi-drug regimen to cure (*1*). One of the key challenges in TB treatment is the complex biology of Mtb bacilli, with heterogeneity in Mtb cell state thought to be a major driver of treatment failure (*2–4*). During infection, Mtb encounters diverse conditions and environmental cues, including acidic pH in the phagosome, lipid-rich environments, and exposure to reactive oxygen species and other immune defenses (*5–8*). As the resident bacteria adapt to this range of conditions, some cells’ growth and metabolic states change, generating phenotypically distinct subpopulations that may survive evading immune clearance and drug killing (*5*, *9–12*). Understanding how drug-tolerant subpopulations of Mtb form in host microenvironments is therefore crucial for improving TB treatment strategies.

One of the earliest stressors Mtb encounters during infection is the acidic pH of the macrophage phagosome (*13*). Before immune activation in macrophages, Mtb inhibits the formation of the phagolysosome and resides in a mildly acidic environment (∼pH 6.2) (*14*, *15*). Mtb adapts to this mildly acidic stress through gene regulation and metabolic shifts that lead to bulk-level growth slowing or growth arrest, which allows Mtb to persist within a host environment (*16–19*). Although these population-level studies provide insights into Mtb’s metabolic plasticity, most studies examine growth rate or growth arrest through bulk measurements, such as optical density (OD_600_) and colony-forming unit (CFU) assays. We therefore lack an understanding of Mtb growth behaviors and changes at the single-cell level during adaptation to host-relevant growth conditions.

Recent advances in single-cell imaging techniques have revealed substantial heterogeneity in mycobacterial growth dynamics, with individual cells exhibiting variations in growth rates, cell cycle timing, and cell size which can affect drug susceptibility (*20–27*). We examined whether growth slowing at acidic pH is uniform across the population or whether there is cell-to-cell variation in growth response. We hypothesized that variability in single-cell responses may be a source of heterogeneity that contributes to such growth slowing and pH adaptation, leading to persistence. Our study revealed that, rather than uniformly slowing growth, Mtb exhibits a bimodal response in which a subset of cells continues to grow at the same rate while another subset converts to a non-growing state. Clinical strains were particularly prone to enter a growth-arrested state. We find that non-growing states are present even in rich, neutral growth conditions, suggesting that Mtb growth arrest is a typical bacterial state and is not due to resource limitations. We challenged Mtb with antibiotic stress and observed that non-growing cells exhibit increased tolerance to the cell wall-targeting antibiotic ethambutol. We identified several regulators involved in modulating this growth behavior under acidic pH conditions, including the two-component system PhoPR, which we also demonstrated is a contributor to growth-arrest-mediated ethambutol tolerance. Our findings suggest that Mtb employs a bet-hedging strategy by changing the proportion of bacteria in a growth-arrested state, with consequences for antibiotic treatment outcome.

## RESULTS

### Growth arrest is the dominant Mtb cell growth state during adaptation to acidic conditions

Many mycobacterial species, including *M. avium*, *M. chelonae*, *M. marinum*, *M. scrofulaceum*, and *M. fortuitum*, grow without restriction under mildly acidic conditions (∼pH 6.0), and some even grow better at acidic compared to neutral pH (*28*). In contrast, Mtb slows its bulk growth rate in response to an acidic environment (*17*, *29*). A simple explanation for the occurrence of growth slowing is that individual cells reduce their growth rates in a uniform manner.

Observations of heterogeneity in mycobacterial growth and metabolic states *in vitro* and *in vivo* led us to speculate that Mtb cells may not uniformly slow their growth (*20*, *21*, *27*, *30–34*). To measure growth changes in single cells as Mtb adapts to acidic pH conditions, we adapted two commonly studied Mtb strains (CDC1551 and H37Rv) and four recently isolated strains of Mtb to pH 5.9 or pH 7.0 conditions for one or four days (Materials and Methods). To identify actively growing cells after different adaptation times, we labeled sites of nascent peptidoglycan within the Mtb cell wall with the fluorescent D-amino acid (FDAA), RADA, in the last 6 hours of the adaptation period (Fig. 1A). The cells were labeled for 6 hours, approximately one-third of the average doubling time, to avoid overcounting growing cells by including those that divide during FDAA treatment. Because mycobacteria elongate from the poles, we can analyze the polar staining patterns of the FDAA to measure growth from each pole over the 6-hour staining period. Consistent with prior studies demonstrating growth heterogeneity in Mtb (*20*, *23*, *27*), we observed variation in the length of the stained polar regions. Specifically, we observed that a large proportion of cells lacked detectable FDAA incorporation, suggesting these cells were entirely non-growing (Fig. 1, A and B). After 4 days of acidic adaptation, the proportion of this non-growing subpopulation among all six strains was 80-95%, whereas the proportion was 34-82% in neutral conditions (Fig. 1B). The bacilli in the non-growing subpopulation were viable, as we observed no difference in survival between Mtb adapted to the acidic or neutral conditions (Fig. S1, p-value = 0.21). These data suggest that even in “fast-growing” conditions, there is a non-negligible subpopulation of non-growing Mtb cells. The proportion of non-growing cells is higher at pH 5.9 than at pH 7.0 for both days of adaptation for all the strains. For example, 59% ± 4.2% of CDC1551 Mtb cells in the neutral condition were growing at day 4, compared with 20% ± 0.2% in the acidic condition (T-test and Benjamini-Hochberg correction p-value = 0.047). Additionally, we observed that the proportion of non-growing cells increased in a pH-dependent manner as shown by comparisons between pH 7.0, 6.2, and 5.9 (Fig. S2). The pattern was clearer at day 1 compared to day 4, likely reflecting differences in bacterial growth phase among the different pH conditions, as cells at neutral pH may have begun entering stationary phase by day 4. We observed a higher proportion of non-growing cells at both pH levels in the clinical isolates compared to the two strains commonly studied in labs (Fig. 1B). This may suggest that having a higher propensity for growth arrest is beneficial for Mtb during infection.

**Figure 1.**
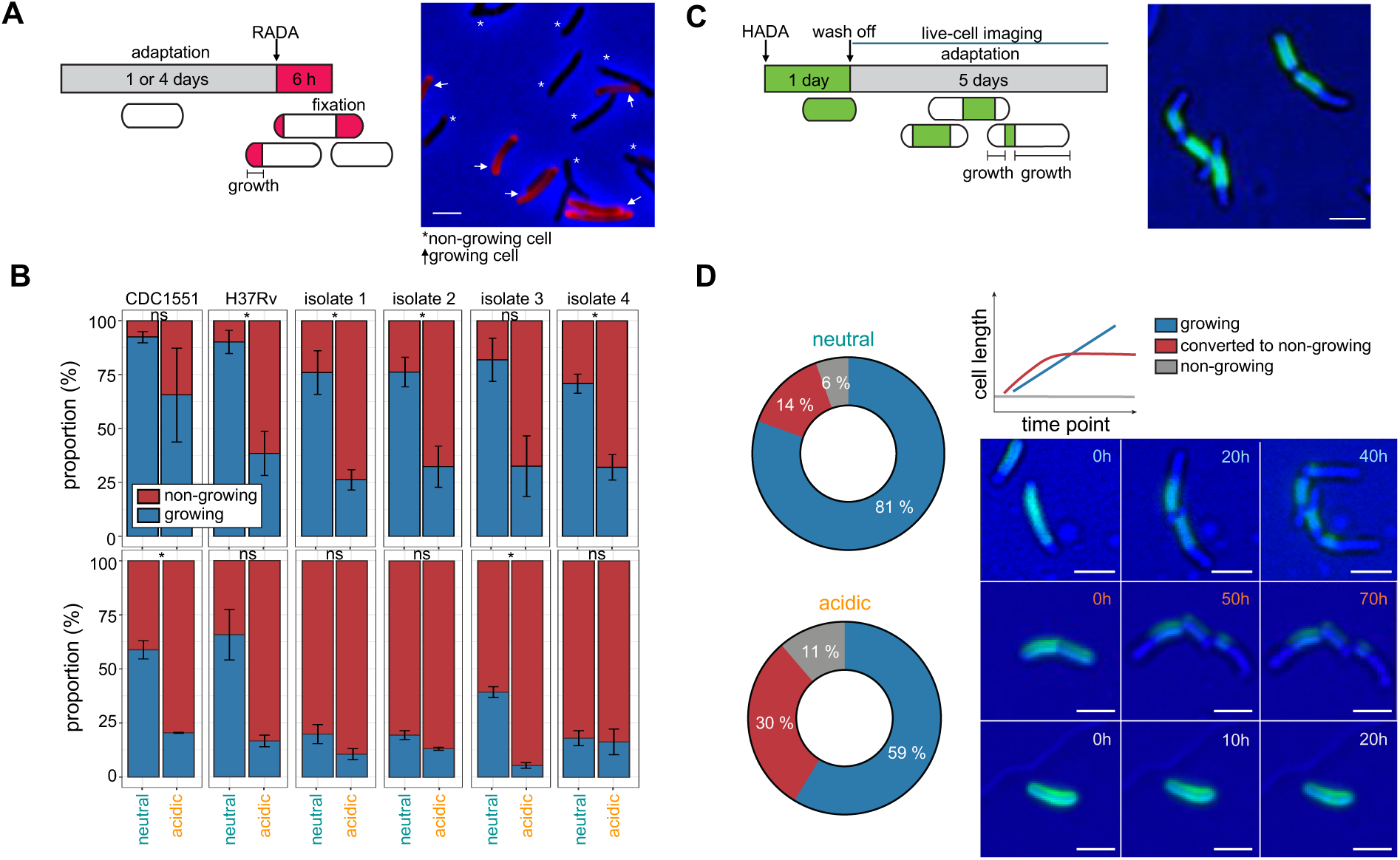
Acidic conditions increase the proportion of non-growing cells. (**A**) Fixed-cell imaging design using fluorescent D-amino acid (FDAA) for growth measurement. RADA was added for 6 hours after 1 or 4 days of adaptation to pH 7.0 or 5.9. Growth is indicated by RADA-labeled length. Scale bar, 2μm. Asterisks and arrows mark non-growing cells and growing cells, respectively. (**B**) Proportions of RADA-positive (growing) and -negative (non-growing) cells in two strains commonly studied in the laboratory (CDC1551, H37Rv) and four recent clinical isolate strains after 1 (top) or 4 (bottom) days of pH adaptation. Significance was determined by unpaired Student’s t-tests with Benjamini-Hochberg correction: ns p ≥ 0.05, *p < 0.05, **p < 0.01, ***p < 0.001, ****p < 0.0001. Data are from biological triplicates. (**C**) Live-cell imaging design using HADA labeling (green). Cells were pre-labeled for 1 day, washed, and then imaged for 5 days under neutral or acidic conditions. New polar growth appears as unlabeled extensions. Scale bar, 2 μm. (**D**) Classification of single-cell growth dynamics during pH adaptation. A schematic diagram categorizes cells into three groups: (1) growing and dividing throughout the movie (green), (2) switched from growing to non-growing (red), or (3) consistently non-growing (gray) throughout pH adaptation (top right). Representative snapshots of each group from live-cell imaging are shown below. A pie chart shows the distribution under neutral (n = 308) and acidic (n = 230) conditions from one experiment. Scale bar, 2μm.

Next, we complemented our fixed-cell imaging assay with a time-lapse pulse-chase imaging experiment to observe the dynamics of the growth adaptation over 5 days (Fig. 1C). We pre-labeled Mtb (CDC1551 strain) with a FDAA (HADA) for one day prior to loading the cells into a microfluidic imaging device with neutral or acidic growth media. We used HADA for the time-lapse imaging because a higher concentration (100 µM) of FDAA was needed to ensure that the whole cell body of each single cell was labeled without disturbing Mtb growth and we found that such a high concentration of RADA disrupted cell elongation. We directly observed growth dynamics by measuring the extension of the cell poles as they elongated with the new (HADA-negative region) growth occurring after the movie started (Fig. 1C, right). We categorized cells tracked over time into three groups based on growth trajectories: “growing,” “non-growing,” and “converted to non-growing” (Fig. 1D upper right). In both neutral and acidic environments, we confirmed that a proportion of cells slowed growth and entered a non-growing state (Fig. 1D, left). Among cells adapted to pH 7.0 (n = 308), 6% were entirely non-growing (n = 17), whereas 14% converted to non-growing (n = 43) during adaptation. These proportions were higher among the cells adapted to pH 5.9 (n = 230), 11% (n = 26) and 30% (n = 69), respectively (Pearson’s Chi-squared test p-value = 2.27×10^-7^) (Fig. 1D, left). Though these proportions do not precisely match the quantities observed by fixed-cell imaging, likely due to differences in assay conditions and methods of growth quantification (see Discussion), they recapitulate the trends in Fig. 1B. The time-lapse and fixed-cell imaging data demonstrate that even in growth-permissive conditions, a fraction of the non-growing population exists, and the proportion of this non-growing subpopulation is higher in an acidic environment.

### The bulk growth slowdown during acidic pH adaptation is caused by an increase in the proportion of a growth-arrested cell subpopulation

Consistent with previous studies, we observed that Mtb growth slows at the population level during adaptation to an acidic environment (Fig. S3A). These data suggest a null model in which the growth of single cells uniformly slows during acidic adaptation, leading to an overall lower average speed compared to growth under neutral conditions (Fig. 2A). However, we observed that there was a growth-arrested cell subpopulation in both neutral and acidic conditions (Fig. 1, B and D), suggesting an alternative model where bulk growth slowdown is caused by an increase in the proportion of cells in the growth-arrested state (Fig. 2B).

**Figure 2.**
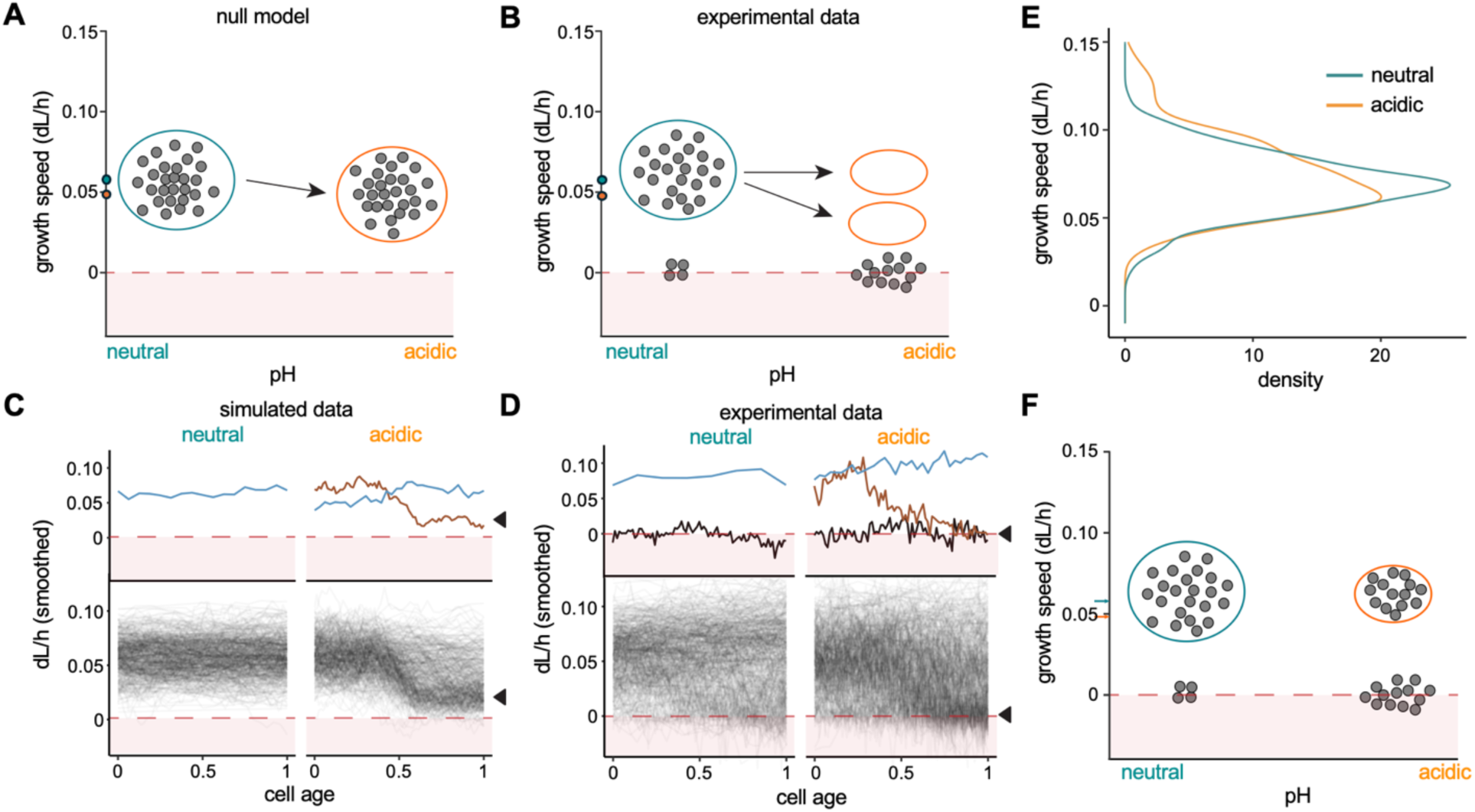
Bulk slowdown of growth during acidic adaptation is caused by an increase in the proportion of growth-arrested cells, rather than a slowed growth speed of growing cells. (**A**) Schematic of null model of single-cell growth rate tuning under neutral and acidic conditions. All cells uniformly slow their growth under acidic conditions. Gray dots represent a single cell. Green and orange circles indicate populations at pH 7.0 and 5.9. Average single-cell growth speeds at pH 7.0 and pH 5.9 are marked on the y-axis. A red dashed line at y = 0 marks growth arrest. (**B)** Schematic of experimentally observed subpopulations of growth-arrested and actively growing cells exist at both pH levels. Growing cells may either maintain (top arrow) or reduce (bottom arrow) their growth rates. **(C–D)** Simulated (**C**) and experimental (**D**) data of single-cell growth speeds (y-axis) over normalized cell age from birth to division. Each line represents a smoothed trajectory of an individual cell using a 20-hour rolling mean, with colored lines highlighting example cell trajectories. (**E**) Density plot comparing cell growth rates (dL/h) of growing cells between neutral and acidic conditions. The Wilcoxon-rank sum p-value = 0.1; n = 248 for pH 7.0, n = 135 for pH 5.9). (**F**) Revised model: the average growth rate decreases under acidic conditions due to an increased proportion of growth-arrested cells, not slower growth in growing cells.

To determine which model explains how Mtb adapts to acid stress, we analyzed single-cell growth rates from our time-lapse imaging experiments (Fig. 1C). We compared these experimental data to time-lapse imaging single-cell data that were simulated under the null model assumptions. According to the null model, we would expect that Mtb in the neutral condition grows at a fairly constant rate above 0 dL/h throughout the 5 days of adaptation (Fig. 2C, left). Instead, in our experimental data, we observed that even in the neutral condition some cells maintained a growth rate of ∼0 dL/h throughout cell-tracking (Fig. 2D, left), confirming that there is a subpopulation of non-growing cells and corroborating our conclusions from Fig. 1. Cells adapting to the acidic condition did not slow their growth rate to above 0 dL/h as in the null model (Fig. 2C, right, arrow); instead, many entered a completely non-growing state at ∼0 dL/h (Fig. 2D, right, arrow).

Additionally, Mtb adapting to acidic pH should slow their growth over the course of tracking time (x-axis), but the final rates should be above 0 dL/h (Fig. 2C, right, arrow). In our experimental data of Mtb adapting to the acidic condition, we observed a subpopulation of cells that continued to grow at a rate similar to the growing cells in the neutral condition, regardless of cell age (Fig. 2D, right). We confirmed this quantitatively, finding that there was no significant difference in growth speeds among cells that continued to grow in neutral and acidic conditions (Fig. 2E, right p = 0.12). These data suggest that Mtb adapts to acidic conditions in a bimodal manner (Fig. S3B), with one subpopulation maintaining the same “fast” growth speed and one growth-arrested subpopulation that increases in abundance (Fig. 2F).

### Non-growing cells that are induced during acidic adaptation recover better from cell wall-targeting drug treatment

To understand whether the growth-arrested subpopulation was differentially susceptible to drug treatment, we measured the drug responses of Mtb adapted to neutral and acidic conditions across a range of antibiotic types (Fig. 3A). Mtb was adapted to either neutral or acidic conditions for four days, then subjected to antibiotic treatment for ten days. After drug removal by transferring the cell cultures onto agar supplemented with 0.4% charcoal (to inactivate antibiotics), the cells were allowed to recover for another ten days. Following recovery, bacterial survival via metabolic activity was assessed by measuring the fluorescence of resazurin. We measured dose responses for four drugs with different mechanisms of action: ethambutol and isoniazid (cell wall-acting drugs), rifampin (a transcription inhibitor), and linezolid (a translation inhibitor) (Fig. 3B, Fig. S4, and Fig. S5).

**Figure 3.**
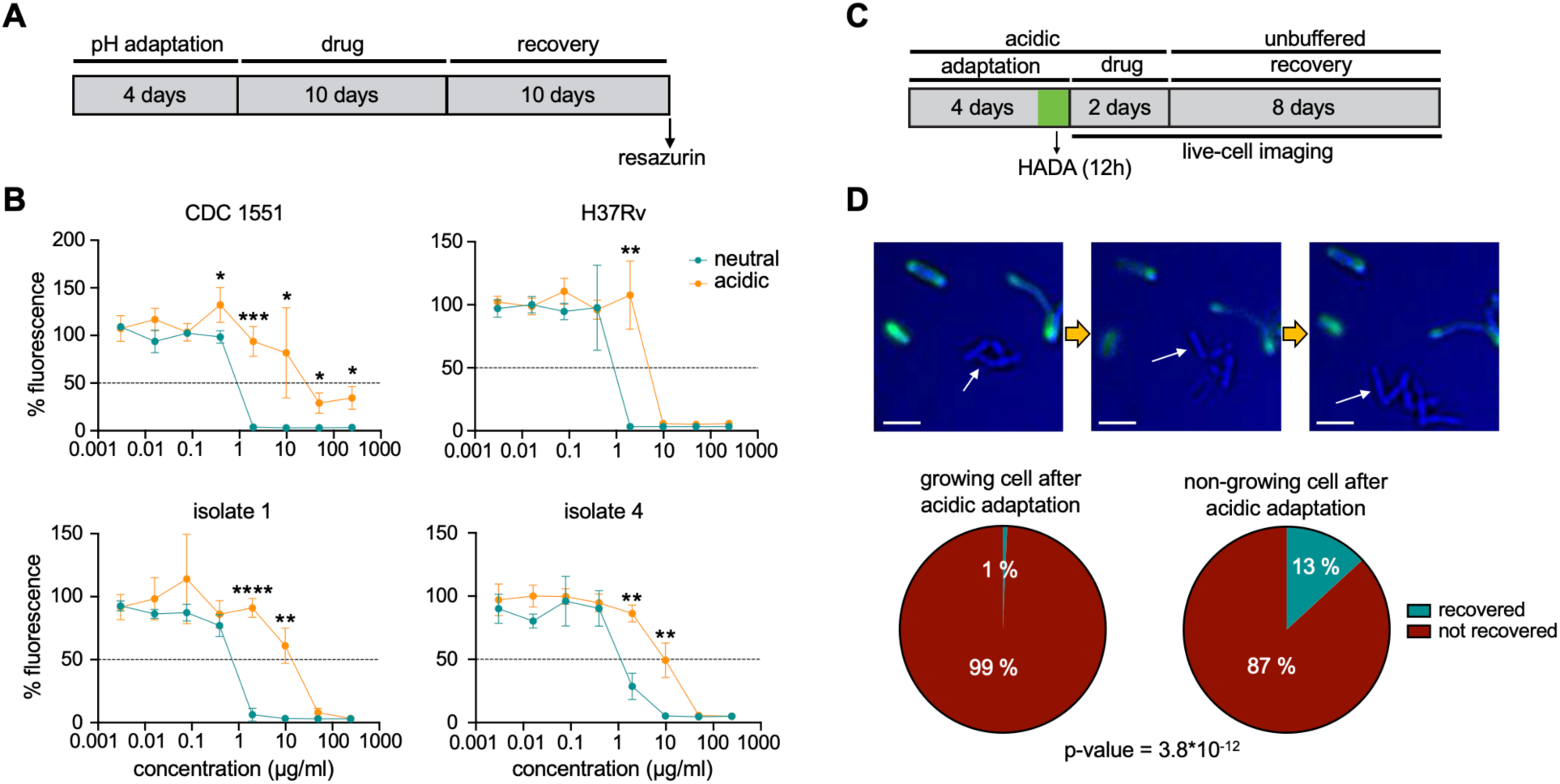
Non-growing cells arising after acidic adaptation recover better from ethambutol than neutral-adapted cells. (**A**) Experimental design for the resazurin assay. Cells adapted to pH 7.0 or 5.9 for 4 days, treated with ethambutol for 10 days, then recovered drug-free for 10 days. Viability was assessed via 1-hour resazurin incubation and fluorescence measurement. (**B**) Four strains (2 lab-adapted, 2 clinical) were tested with varying ethambutol concentrations. Green = neutral-adapted; orange = acidic-adapted. Fluorescence was normalized to untreated controls. Fluorescence level of resazurin reflects metabolic activity of bacterial cells. Mean ± SEM shown. P values were calculated using the two-tailed unpaired t-test. (**C**) Experimental design for drug treatment time-lapse imaging. CDC1551 WT was adapted to pH 5.9 for 4 days. HADA was added for the last 12 hours. Cells were imaged for 10 days, with 245 µg/ml ethambutol for the first 2 days, followed by fresh media. (**D**) Growth resumption was assessed in HADA-positive (growing, *n* = 736) and HADA-negative (non-growing, *n* = 152) cells. Fisher’s exact test was used. Arrows highlight previously non-growing cells resuming growth.

When Mtb were adapted to acidic conditions, they exhibited significantly lower susceptibility to ethambutol compared to when they were adapted to neutral conditions across all the strains tested (Fig. 3B). At 2 µg/ml ethambutol, all four strains in neutral conditions exhibited a drastic reduction in survival, with no recovery observed at 10 µg/ml. Significant differences were detected between neutral and acidic adaptation at both concentrations: at 2 µg/ml, p-values were 0.0006, 0.003, 8.8×10^-5^, and 0.001 for each strain, respectively; at 10 µg/ml, p-values were 0.05, 0.002, and 0.005, excluding the H37Rv strain, respectively (Fig. 3B). In contrast, acidic-adapted cells demonstrated robust recovery at 2 µg/ml ethambutol and maintained partial recovery even at 10 µg/ml, a concentration at which neutral-adapted strains failed to recover.

Across the Mtb strains, we did not observe consistent differences in susceptibility to isoniazid, rifampicin, or linezolid following acidic pH adaptations (Fig. S4 and S5), in contrast to the uniform response seen with ethambutol (Fig. S4 and S5). Rifampicin and linezolid are known to be effective against bacteria in non-replicating states; therefore, we did not expect that an increased proportion of growth-arrested (e.g., non-replicating) bacteria resulting from acidic adaptation would decrease susceptibility to these drugs. Acidic adaptation may not impact isoniazid treatment in the same way as ethambutol because acidic adaptation is thought to increase inhibition of the target of isoniazid, InhA, via PhoPR-dependent upregulation of NAD⁺ (*17*, *29*), thereby increasing isoniazid susceptibility via a mechanism independent of growth arrest. These findings suggest that acidic adaptation enhances Mtb tolerance to ethambutol, but not isoniazid, rifampicin, or linezolid.

The resazurin assay reports bulk-level metabolic states as a measure of survival to drug treatment, these dose-response assays do not distinguish between the relative survival of cells that were in the growing versus the growth-arrested subpopulations. We hypothesized that ethambutol tolerance arises primarily from the non-growing cell subpopulation because ethambutol inhibits arabinogalactan synthesis, a process essential for cell wall elongation in actively growing cells (*35*, *36*). To test this hypothesis, we conducted live-cell imaging to assess which cells (growing or non-growing) resumed growth after drug removal (Fig. 3, C and D). To identify individual cells as growing or non-growing immediately preceding drug exposure, the CDC1551 WT strain was adapted to pH 5.9 for four days and labeled with HADA. Cells were then loaded into a microfluidic device for time-lapse imaging, where they were treated with a high ethambutol dose (245 µg/ml) for two days and allowed to recover for eight days in drug-free fresh media. This high ethambutol dose was required in live-cell imaging due to differences in environmental conditions compared to the fixed-cell assay, such as the continuous flow of fresh, nutrient-rich media through the cells and the high air permeability of the device (details in Materials and Methods).

We observed that non-growing cells formed during acidic adaptation predominantly contributed to recovery after the ethambutol treatment (Fig. 3D and Table S1). Only 1% of previously growing cells resumed growth upon ethambutol removal (5 out of 736 cells), whereas 13% of the non-growing cells recovered (20 out of 152 cells) (Table S1). Together, these data suggest that non-growing cells that arise after acidic adaptation have reduced ethambutol susceptibility.

### PhoPR is a partial regulator of growth and ethambutol tolerance in acidic adaptation

PhoPR is a two-component regulatory system and a key regulator of cellular adaptation to acidic conditions (*29*, *37*, *38*). We hypothesized that PhoPR controls the proportion of cells in the non-growing subpopulation and is therefore required for ethambutol tolerance attributed to growth arrest from adaptation to acidic conditions. To assess the extent to which PhoPR regulates growth arrest, we performed fixed-cell imaging using FDAA labeling in CDC1551 WT, *phoPR*-deleted mutant (Δ*phoPR)*, and *phoPR*-complemented (*phoPR*\*) strains at neutral (pH 7.0) and acidic conditions (pH 5.9 and 5.7) (Fig. 4A). Though increased non-growing cell formation was observed at pH 5.9 compared to pH 7.0, the result was independent of PhoPR, as non-growing cell proportions did not differ between WT and *phoPR*\* strains compared to the Δ*phoPR* strain. However, at pH 5.7, *phoPR*-dependent non-growing cell formation was evident, with significantly fewer non-growing cells in the Δ*phoPR* strain than in WT and complemented strains (p-value 0.004 and 0.009, respectively). These data suggest that the PhoPR regulon is partially involved in increasing the proportion of the growth-arrested subpopulation at pH 5.7.

**Figure 4.**
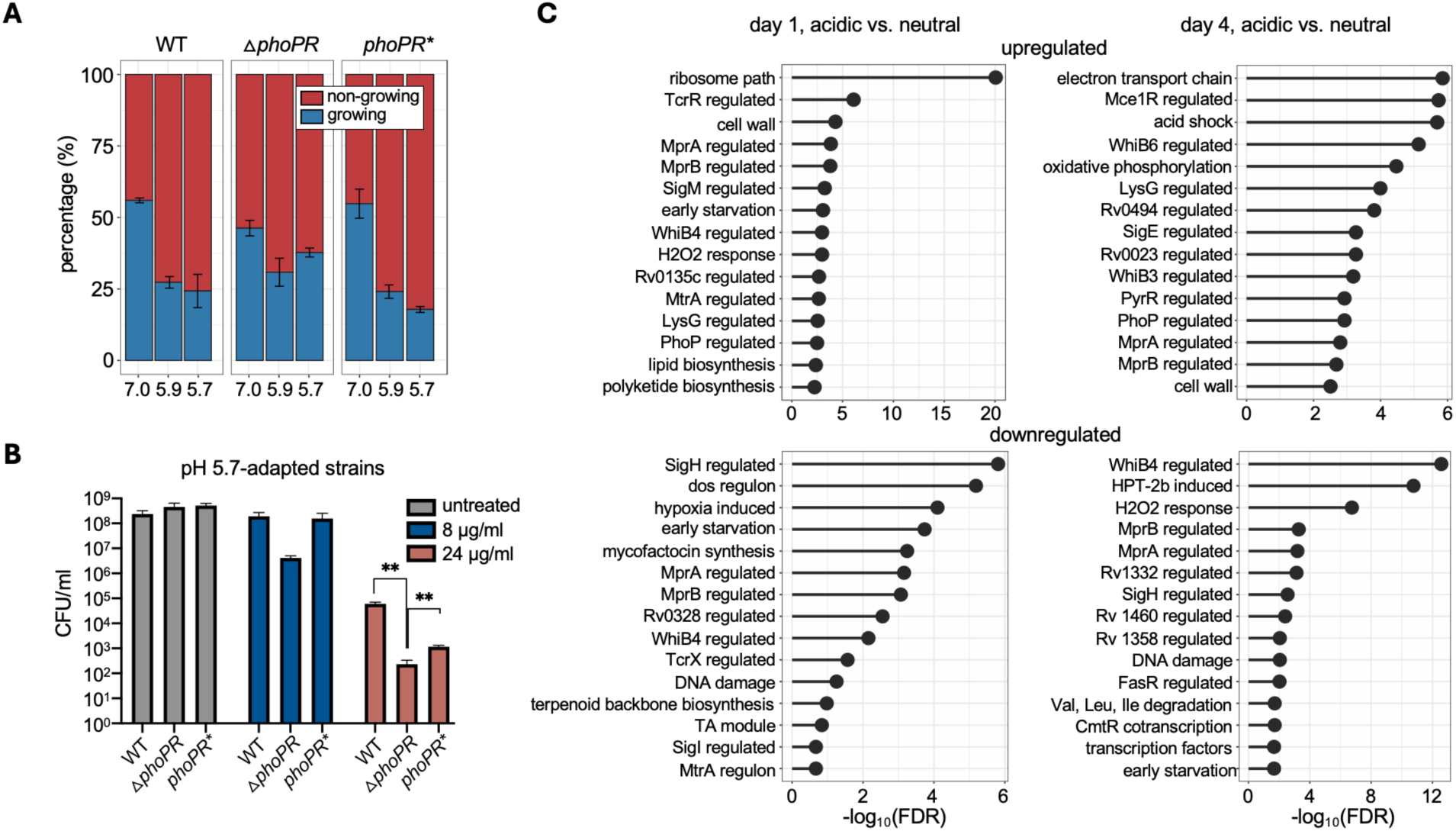
PhoPR is involved in growth arrest and ethambutol susceptibility under acidic conditions. (**A**) Proportions of growing (HADA-positive, blue) and non-growing (HADA-negative, red) cells after 4-day adaptation to pH 7.0, 5.9, or 5.7. Three strains were used – CDC1551 streptomycin-resistant WT (STR-R), *phoPR* mutant, and complemented strains (details in Materials and Methods). See Fig. 1A for experimental design. (**B**) After a 4-day adaptation at pH 5.7, cells were treated with ethambutol (untreated, 8, or 24 µg/ml) for ten days, then plated for CFU. Biological triplicates; mean with SEM. p-values between WT and mutant (p = 0.004), and mutant and complemented (p = 0.009); unpaired two-tailed t-test. (**C**) Multi-strain pathway and gene regulator enrichment analysis from RNA-seq of CDC1551, isolate 1, and isolate 2 after one or four days at neutral and acidic (pH 5.9). Differentially expressed genes (DEGs) (adjusted p-values < 0.05), shared across all strains under acidic conditions, were analyzed to identify commonly regulated pathways by the upstream regulators (hypergeometric test). The readout represents the statistical significance of differential gene expression. FDR, false discovery rate.

We examined the role of PhoPR in ethambutol tolerance under acidic conditions by comparing survival among the WT, Δ*phoPR* mutant, and *phoPR\** strains. Survival rate was measured by counting colony forming units (CFU) at pH 5.7 using these three strains and two ethambutol concentrations derived from Fig. 3B (Fig. 4B). The two ethambutol doses were chosen to capture the dose where survival of acidic-adapted cells began to decrease, allowing for comparison of drug tolerance across strains. In the absence of ethambutol, a similar CFU/ml was observed among all strains. However, upon treatment with 24 µg/ml ethambutol, the Δ*phoPR* mutant exhibited significantly lower CFU counts than WT and *phoPR\** strains, suggesting that PhoPR contributes to the ethambutol tolerance observed in this context. Although a similar trend was observed at 8 µg/ml ethambutol, the difference was not statistically significant.

Because deletion of *phoPR* can influence cell wall lipid composition (*39*), we examined whether the increased ethambutol susceptibility of Δ*phoPR* was due to higher drug permeability compared to WT. We found that Δ*phoPR* was more permeable to the fluorescent stain, FM 4-64, than WT and *phoPR\** at pH 5.7 and at pH 7 (Fig. S6). Because there was no difference in ethambutol susceptibility between WT and Δ*phoPR* at pH 7.0 (Fig. S6), we conclude that the increased permeability in Δ*phoPR* does not lead to increased ethambutol susceptibility. Together, these data suggest that the Δ*phoPR* Mtb is more sensitive to ethambutol than WT Mtb, not because of increased drug permeability, but because of an increase in the proportion of non-growing cells.

### Multiple regulatory pathways are involved in Mtb acidic adaptation

Collectively, these findings demonstrate that although PhoPR is a key regulator of growth arrest and drug tolerance under acidic conditions (in pH 5.7), additional regulatory mechanisms are likely involved in this complex adaptation. To identify other possible regulatory mechanisms involved in the transition to growth arrest and ethambutol tolerance under acidic conditions compared to neutral conditions, we performed RNA sequencing (RNA-seq) on CDC1551 WT and two clinical isolates after four days of adaptation to neutral (pH 7.0) or acidic (pH 5.9) conditions. To determine pH-specific transcriptional changes common to all strains, we took the intersection of their differentially expressed genes and subjected them to pathway and gene regulatory program enrichment analysis. We found that PhoP-regulated genes were one of the most differentially expressed genes in acidic conditions across all tested strains (Fig. 4C, adjusted p-value = 3.1×10^-3^). However, there were other regulatory pathways that were also upregulated, suggesting that while PhoPR plays a key role, it may not be the sole master regulator of growth control in acidic environments. Specifically, other transcriptional programs that are known to modulate growth and division and were more enriched than the PhoP-regulated gene sets included MprAB-, WhiB4-, WhiB3-, MtrA-, TcrX-, and SigH-regulated pathways (Fig. 4C). Of note, the genes regulated by the MprAB two-component system were significantly upregulated at both day 1 and day 4 of adaptation (mean adjusted p-value = 1.3×10^4^ and 1.83×10^3^, Fig. 4C). MprAB responds to pH and cell envelope stress and coordinates the expression of over 200 genes that control resistance to cell envelope damage and growth speed. The MprAB system may thus play a key role in generating a non-growing cell subpopulation in acidic conditions that is tolerant to ethambutol-induced cell wall damage.

## DISCUSSION

The ability of Mtb to adapt to different environments during infection, including to acidic pH, is critical to Mtb survival in the host and to drug treatment, yet we lack an understanding of how individual cells adapt to these environments. Here, using single-cell approaches, we find that individual bacteria adapt to acidic pH conditions in a manner that could not be inferred from population-level measurement alone. Instead, we find that Mtb adapts to acidic pH not by uniformly slowing growth, but by increasing the proportion of cells in the non-growing subpopulation. We observed a significant proportion of non-growing cells not only under acidic pH conditions but also in adaptation to neutral conditions. These data suggest that non-growth is not a rare state but rather a common state for Mtb even in nutrient-rich conditions with limited environmental stress. The propensity to enter this non-growing state was higher in recently isolated clinical strains compared to those commonly studied in the laboratory. Clinical isolates have had fewer passages in laboratory growth conditions. Therefore, we expect that they would retain adaptations from the human host, which would contribute to the phenotypic plasticity. Mtb are known to enter non-growing states under severe stressors, such as hypoxia, or in lipid-rich conditions of the caseum. But our data demonstrate that there is an abundance of non-growing cells emerging after short adaptations even to rich, mild conditions in the laboratory, suggesting that a growth-arrested state may be more typical for Mtb than previously thought.

We found that the bulk slowed growth at acidic pH is due to an increased proportion of the non-growing cell subpopulation, not to slower single-cell growth rates (Fig. 2C, 2D, and Fig. S3). Correspondingly, cells that continued to grow throughout the experiment in the acidic condition maintained the same growth rate as those in the neutral condition (Fig. 2E). These data show that Mtb tunes its growth rate not by uniformly slowing or accelerating single-cell growth speeds but instead by tuning the proportion of growth-arrested cells in the population, i.e., cells do not slightly change their growth rates at the single-cell level but rather shift from a regular growth mode to a growth-arrested mode. Maintaining this heterogeneous population of cells likely provides a survival benefit in the face of various stressors (bet-hedging) (*10*, *40–42*).

We found that the growth-arrested subpopulation in the acidic condition was more tolerant to ethambutol treatment than the growing subpopulation (Fig. 3, B and D). Antibiotic efficacy is known to be strongly dependent on the physiologic state of the bacteria (*43*), and previous studies in mycobacteria have confirmed that this is true on the single-cell level (*20*, *21*, *23*, *44*). In this study, the acid-adapted Mtb that were growth arrested were protected against ethambutol across different strains. We note that protection of the growth-arrested subpopulation did not generalize to other antibiotics tested (Fig. S4 and Fig. S5), demonstrating that the physiologic state of the non-growing cells may be protective against some but not all types of stressors. Future studies may examine if growing and non-growing subpopulations in acidic conditions are differentially susceptible to host-relevant stresses, such as reactive nitrogen species, reactive oxygen species, or hypoxia.

We did not examine the mechanisms by which Mtb exposed to acid stress better survives ethambutol treatment, but it is possible that changes in cell wall-related gene expression play a major role. A gene *embB*, which encodes a protein involved in the biosynthesis of arabinogalactan of the cell wall, mediates resistance to ethambutol treatment (*45–47*) and was significantly upregulated in acidic conditions compared to neutral conditions in the three tested strains, which included two clinical strains (Table S2). Future studies may examine whether the upregulation of *embB* directly contributes to the higher survival of Mtb in acidic conditions.

Although both isoniazid and ethambutol target the mycobacterial cell wall, their effects on bacterial survival after acidic adaptation diverged due to distinct mechanisms of action. Isoniazid is a prodrug that is activated by the catalase-peroxidase KatG, producing isonicotinoyl radicals that bind NAD⁺ to form isonicotinoyl-NAD (*48*). Isonicotinoyl-NAD inhibits InhA, a key enzyme involved in mycolic acid biosynthesis, thereby compromising cell wall integrity (*49*, *50*). Under isoniazid pressure, Mtb downregulates genes encoding NADH dehydrogenase to limit NAD⁺ production, reducing isonicotinoyl-NAD formation and thus mitigating InhA inhibition to promote survival (*51–53*). However, under acidic conditions, PhoPR regulation enhances NAD⁺ production, which results in increased isonicotinoyl-NAD formation, increasing bacterial susceptibility to isoniazid through enhanced InhA inhibition (*54–56*). These findings underscore that drugs with similar targets, such as ethambutol and isoniazid, can yield distinct outcomes under host-relevant stress conditions due to differences in their activation pathways.

PhoPR is a master regulator of Mtb’s transcriptional response to acidic conditions (*13*, *17*, *29*, *37*, *57*), but we found that it was only partially responsible for controlling the proportion of the non-growing cell subpopulation in response to pH fluctuation (Fig. 4A). To further explore additional regulatory systems beyond PhoPR, we examined transcriptional changes associated with acidic adaptation across three strains. The most significantly upregulated pathway between neutral and acidic conditions common to all strains tested was the ribosome biogenesis, potentially reflecting a compensatory mechanism to counteract reduced protein synthesis rates caused by acid stress (*58*, *59*). While PhoP-regulated genes were significantly differentially expressed, other regulators involved in sensing pH, altering growth and division, and resisting cell wall stress were more strongly enriched (Fig. 4C). For example, MprAB is known to respond to pH and controls the expression of genes that provide resistance against cell wall damage. The MtrAB two-component system is also known to play a crucial role in dormancy, cell division, cell wall metabolism, and modulating drug susceptibility and MtrA-regulated genes are significantly differentially expressed at acidic pH (*60–65*). More comprehensive genetic analysis may find that MprAB, WhiB4, SigH, or other differentially expressed regulons play a larger role than PhoPR in generating the non-growing cell subpopulation. It may be that many transcriptional regulators are involved to provide redundancy and robustness to the acid stress response in Mtb. It is also likely that some regulators respond to multiple cues to integrate physiological responses; for example, Mce3R integrates cues from carbon sources and potassium (*66*). Simultaneous integration of multiple environmental cues may allow Mtb to fine-tune its transcriptional response. Another explanation for why so many transcriptional regulators are involved in responding to acid stress could be that early and delayed responses requires different regulation. In support of this, genes regulated by the redox-dependent WhiB4 regulator are upregulated at day one but downregulated by day four (Fig. 4C) (*67*).

In addition to transcriptional regulation, carbon source availability can drive metabolic shifts that influence bulk slow growth or growth arrest under acidic conditions (*16–19*). For example, Mtb fully arrests growth at mildly acidic pH (pH 5.7) when provided with glucose, glycerol, or most TCA cycle intermediates as sole carbon sources (*18*). In contrast, Mtb can maintain growth at pH 5.7 when utilizing alternative carbon sources such as cholesterol, acetate, and oxaloacetate (*18*, *19*, *66*). In this study, Mtb strains were grown in 7H9 medium supplemented with OADC (oleic acid, albumin, dextrose, catalase), a rich carbon source mixture. Therefore, a limitation of our study is that we did not assess growth patterns under conditions using individual carbon sources.

Many pathogens, such as *Streptococcus, Escherichia, Salmonella*, and *Pseudomonas*, have adapted mechanisms to survive at acidic pH. These include the activation of acid resistance systems and the downregulation of H^+^ pumps to limit proton entry into the cell (*68*, *69*). In macrophage environments, subpopulations of *Salmonella* are known to slow their growth while maintaining an active metabolism (*70*). It may be that Mtb and *Salmonella* similarly induce growth arrest in a subpopulation while another subpopulation continues to grow at the same rate as a bet-hedging strategy (*70*). Future studies may investigate whether other pathogens coordinate bifurcations of growth and how this mechanism impacts bacterial fitness.

Together, our study highlights how single-cell approaches can provide key insights that may be overlooked by population-level measurements alone. By understanding behaviors and variation of individual cells, we can investigate how different subpopulations arise and respond to host- and environmental stressors. Future studies may also aim to identify the genetic and metabolic regulators that control the growth-state decisions in individual cells, as these factors may serve as critical targets for therapeutic intervention.

## MATERIALS AND METHODS

### Mtb strains

Six Mtb strains were used in this study: CDC1551 and H37Rv and four clinical strains, 24TB069, 24TB041, 24TB25, and DTU117 from (*22*) that are referred to as isolates 1 through 4, respectively. These strains were selected to reflect the diversity of Mtb genetics and virulence present in TB patients in Vietnam. The strains encompass all three major Mtb lineages in TB patients in Vietnam: 24TB069, Indo-Oceanic; 24TB041, East-Asian; 24TB25, East-Asian; and DTU117, Euro-American. They were also selected to represent a range in degrees of virulence in a macrophage lysis model: 24TB069, moderate; 24TB041, high; 24TB25, high; and DTU117, high. The CDC1551 Δ*phoPR*, and *phoPR\** (complemented mutant) strains have been previously described (*37*).

### Bacterial culture

Mtb strains were grown in standard medium consisting of 7H9 broth (ThermoFisher; DF0713-17-9) with 0.05% Tween 80 (ThermoFisher; BP338-500), 0.2% glycerol (ThermoFisher; G33-1), and 10% Middlebrook OADC (ThermoFisher; B12351). Strains were grown to OD_600_ of 0.5-1.0 from frozen at 37°C with mild agitation (100 rpm). They were back-diluted to OD_600_ 0.05 and grown to mid-log phase (OD_600_ 0.5-1.0) before experimental use.

#### For fixed-cell imaging

Strains were passaged to OD_600_ 0.05 in three different media types: standard 7H9 pH 7.0, pH 6.2, and pH 5.9. pH-adjusted 7H9 media were made with 100mM MOPS buffer (SigmaAldrich; M3183) added to standard medium and NaOH to adjust pH to 7.0, or 100mM MES buffer (SigmaAldrich; M2933) added to standard medium and NaOH to adjust the pH to 6.2 or 5.9. Each culture was placed on 96-well plates (150 µl/well) and incubated at 37°C in humidified bags until fixation. Three biological replicates were generated for each strain and each condition.

#### For live-cell imaging

Strain CDC1551 was used for live-cell imaging. Prior to live-cell imaging experiments, Mtb cells were stained with 100 µM of HADA in 10ml standard 7H9 broth media for 24 hours. Cells were then washed twice with phosphate buffered saline with 0.2% Tween-80 (PBST) and resuspended in pH 7.0 or pH 5.9-adjusted 7H9 media.

### Fluorescent staining

The fluorescent D-amino acids (FDAA) RADA (Tocris; 6649) and HADA (Tocris; 6647) were used in this study. FDAA powder was dissolved in DMSO to a stock concentration of 100 mM and stored short-term at −80°C. Cells were incubated in 20µM RADA or HADA for 6 h before fixation at each time point. For live-cell imaging, cells were incubated in 100 µM HADA for either 12 hours or 24 hours prior to the start of the imaging. For the permeability assay, fixed Mtb were stained in a solution of 5.2 µg/mL of FM4-64FX (ThermoFisher; F34653), PBS (ThermoFisher; 20012-027), and PBST. Cells were incubated in the presence of dye at room temperature in the dark for 30 minutes. Stained cells were washed once with 150 µL of PBST prior to imaging.

### Fixation of Mtb cells

At designated time points, Mtb cultures were fixed in paraformaldehyde (Alfa Aesar, 43368) at a final concentration of 4% for one hour, and removed from the biosafety level-3 facility. After fixation, the samples were washed twice with PBST and resuspended in PBST. Plates were sealed (ThermoFisher optically clear plate seals; AB1170), and all samples were stored at 4°C until imaging.

### Fixed-cell imaging

Fixed Mtb cells were spotted onto 1% agarose pads (SigmaAldrich; A3643-25G) as in (*71*). In Fig. 1, images were captured with a widefield DeltaVision PersonalDV (Applied Precisions) microscope. Bacteria were illuminated using an InsightSSI Solid State Illumination system with transmitted light for brightfield microscopy. RADA was visualized with Ex. 544nm and Em. 570nm wavelengths. Montage images were generated using a custom macro that captures 25 individual fields of view per image. Images were recorded with a DV Elite CMOS camera for all channels. For imaging of the WT, Δ*phoPR*, and *phoPR\** strains (Fig. 4), a widefield Zeiss Axio Observer 7 with Definite Focus microscope and Hamamatsu ORCA-Fusion Digital CMOS camera was used. Bacteria were illuminated with phase contrast and HADA was visualized with Ex. 370-400 nm and Em. 410-440 nm using the 90 HE DAPI reflector and the LED-Module 385 nm light source. FM 4-64 stained Mtb were imaged with Ex. 489-533 nm and Em. 542-832 nm using the 91 HE filter set and the LED-Module 511 nm light source.

### Fixed-cell image segmentation

Before image segmentation, ImageJ plugin BaSiC was used to ensure an even distribution of illumination in all channels across the image (*72*). Image segmentation and feature extraction were performed using a custom pipeline that relies on ilastik (v1.4.0) and CellProfiler (v4.2.1). Briefly, an ilastik pixel classifier was trained to distinguish between cell and background pixels and applied to all images. Using these pixel prediction maps, an ilastik object classifier was trained to discriminate single cells from clumps of cells or debris in the images. FDAA-positive regions were segmented using a separate ilastik pixel classifier trained to distinguish between fluorescent pixels and background pixels. The resulting cell boundary predictions and regions of FDAA positivity were then used as input for a CellProfiler pipeline. These two ilastik prediction sets were converted to objects in CellProfiler using the “IdentifyPrimaryObjects” module with manual thresholding, and the “RelateObjects” module was used to relate the FDAA-positive regions to each segmented cell. The single-cell measurements were exported in CSV format using the “ExportToSpreadsheet” module for downstream analysis and plotting.

### Live-cell microscopy

Before loading Mtb cells into a custom polydimethylsiloxane (PDMS) microfluidic device, cells were filtered through a 10 µM membrane filter to remove clumps. Mtb cells were then loaded into a microfluidic device, as in (*21*). The device contained a main microfluidic feeding channel with a height of 10-17 µm and viewing chambers with a diameter of 60 µm and a height of 0.8-0.9 µm. Fresh medium was delivered to the cells at 5 µl/min flow using a microfluidic syringe pump. The device was placed on an automated microscope stage within an environmental chamber maintained at 37°C. Mtb cells were imaged for 120 hours at one-hour intervals using a widefield Deltavision PersonalDV (Applied Precision) microscope. Cells were illuminated with an InsightSSI Solid State Illumination system every hour. Cells were imaged using transmitted light brightfield microscopy, and HADA was visualized with 425-445nm excitation and 460-510nm emission wavelengths.

#### Live-cell imaging with ethambutol

For ethambutol-treated live-cell imaging, the CDC1551 WT strain was used. Prior to imaging, the cells were pre-adapted to pH 5.9 7H9 media for four days. The HADA (100 µM) was added during the last 12 hours of the adaptation period to mark whether the cells were growing (HADA positive) or non-growing (HADA negative) after adaptation. For the pH 5.9 media that was flowed in the microfluidic device, media conditioned using TB auxotroph strain adapted to pH 5.9 7H9 media was mixed 1:1 with pH-adjusted regular 7H9 media (*27*, *73*). Once the cells were loaded into the device, the pH-adjusted media containing 245 µg/ml of ethambutol was delivered through the device. After two days, the syringe containing ethambutol was replaced with a new syringe containing fresh, unbuffered media without ethambutol for the recovery period. For this unbuffered media, unbuffered spent media was mixed 1:1 with regular unbuffered 7H9 media. The recovery period was imaged for eight days. The cells were imaged with bright field (32% transmission, 0.1 s exposure time) and CFP filter (5% transmission, 0.1 s exposure time).

### Live-cell image annotation

Before segmentation, each channel was merged into one image. ImageJ plugin ObjectJ was used to hand-annotate cell length and growth throughout live-cell imaging. Each annotated movie was then extracted as a CSV file for further analysis.

### Segmented data analysis

Among the annotated cells, those which could not be determined to be either growing cells or cells that converted from growing to non-growing were excluded from the analysis. This includes cells that were born late in the movie or cells that could no longer be annotated due to clump formation or overlap with other cells due to high cell density. The annotation in each frame was extracted, containing information on cell length and growth at each pole over time (1 hour timescale). The ObjectJ data were exported to an XML file, then converted to a CSV file. For the single-cell analysis, we used custom scripts in MATLAB that calculated and collated single-cell data - length at birth and division, growth from each pole, and interdivision time. To classify cells as non-growing, growing, and growing to non-growing, we used a combination of quantitative and visual annotation. First, we smoothed hourly growth rates using a lowpass filter with coefficients equal to 5 using the “smooth” function in MATLAB. Cells whose smoothed growth rate throughout the movie was close to 0 dL/dT were classified as non-growing. Cells whose smoothed growth rate was > 0/dL/dT and fitted a linear regression with low error, R-squared > 0.9, were classified as growing. If the fit was < 0.9, then the cells were classified as “growing to non-growing,” since a poor fit indicates the cell growth did not follow a linear trajectory and instead must have slowed. Accurate binning into these categories was confirmed by plotting cell length over time and visually inspecting the slopes.

### Growth rate analysis from live-cell imaging

To compute the instantaneous growth rate of Mtb cells, we measured cell lengths hourly over the course of the experiment. To account for cell length variations caused by microscopy field of view instability, we computed a smoothed instantaneous growth rate using a 20-hour rolling window. For the simulated null model data, instantaneous growth rates were generated by replacing the experimental data with randomly sampled values so that the final population-averaged growth rate matched that of the experimental data. For the neutral pH simulated data, these values were sampled from a distribution with a mean equal to that of the neutral pH experimental growth rate, mean = 0.058 dL/h, and S.D. = 0.08 dL/h. For the acidic pH simulated data, we inputted values sampled from a distribution equal to the neutral pH simulated values (mean = 0.058 dL/h and S.D. = 0.08 dL/h) for the first 48 hours. After 48 hours of experiment time, the values were randomly sampled from a distribution of mean = 0.04 dL/h and S.D. = 0.02 dL/h, to reflect the null hypothesis that single-cell growth should be slowing but not halting. Again, these values were chosen so that the acidic pH simulated data’s population-averaged growth rate matched the 0.049 dL/h mean of the experimental data. The resulting dL/h measurements were smoothed with a 20-hour rolling window in the same way as the experimental data.

### Preparation of charcoal agar plates

Charcoal agar plates were prepared as previously described (*74*). Briefly, 450 ml of Middlebrook 7H10 agar containing 0.2% glycerol and 0.4% of activated charcoal was autoclaved in a flask. Once the autoclaved agar cooled down to 55-65°C, the flask was placed on a magnetic stirrer hot plate to maintain the charcoal in suspension. Then, 10% Middlebrook OADC (ThermoFisher; B12351) was added. After pouring the 7H10-OADC-charcoal into a reservoir, 200 µl of 7H10-OADC-charcoal was transferred to each well of a 96-well plate using a 12-multichannel pipette. Once the poured agar solidified, the plates were placed in plastic bags and stored at 4°C until use.

### Resazurin assay after antibiotic treatment

Strains were adapted to their respective pH-buffered media for four days prior to antibiotic treatment. Cells were then back-diluted to OD_600_ 0.05 and treated with drugs using various concentrations in 384-well flat clear bottom plates. Cells without drugs were used as a control. After ten days of drug treatment at 37°C incubation, 10 µl of the cell culture was transferred into a new 96-well plate containing 200 µl of 7H10-OADC-charcoal per well to allow the charcoal to sequester the drugs used to treat Mtb (*75*). After 10 days of recovery, 40 µl of sterile 1xPBST (0.05% Tween-80 in PBS) was added to prevent the dried charcoal from absorbing resazurin. 50 µl of freshly prepared resazurin solution – 0.01% of resazurin in 5% Tween-80 in PBS, filtered – was added to each well. Then the plates were rocked back and forth a few times, placed in a plastic bag and a secondary container, and incubated at 37°C for one hour. The fluorescence was read using a Biotek Synergy Neo2 Hybrid Multi-Mode Reader, with excitation at 530 nm and emission at 590 nm.

### CFU assay

#### CFU assay after pH adaptation

The CDC1551 WT strain was used for the CFU assay after pH adaptation to evaluate viability after acidic adaptation. The strains were adapted to either pH 7.0 or pH 5.9 for four days, followed by serial dilution and plating on 7H10 agar plates. The colony was counted after three weeks. The assay was performed with biological triplicates.

#### CFU assay for drug treatment

CDC1551 WT (STR-R), Δ*phoPR*, and *phoPR\** strains were used for the assay. Strains were first adapted to either pH 7.0 or pH 5.9 for four days prior to antibiotic treatment. After four days of pH adaptation, cells were back-diluted to OD_600_ 0.05 using each pH-adjusted medium and treated with ethambutol at 8 or 24 µg/ml. Untreated samples were used as a control. After ten days of ethambutol treatment, cells were washed twice with unbuffered 7H9 media for 10 minutes at 2,500 rpm. The cells were resuspended in unbuffered 7H9 media, serially diluted, and plated on 7H10 agar plates. The colony counting was performed after three weeks. The assay was performed using biological triplicate.

### Sample preparation for RNA sequencing

Biological triplicates of CDC1551, V1, and V3 strains were used for RNA sequencing. When the cells were at mid-log phase (OD_600_ 0.3-0.5) in the unbuffered media, the cells were back-diluted to OD_600_ 0.05 in 10ml of buffered media (pH 7.0 or pH 5.9). The cells went through adaptation to each pH-adjusted medium, and the cells were then harvested by centrifugation at two different time points, after one or four days of adaptation. At each time point, GTC buffer was added to halt transcriptional response during RNA extraction, and the cells were washed twice. The cell pellets were resuspended in ∼1ml of GTC buffer (4M GTC + 0.5% sarcosyl + 25 mM sodium citrate + 0.1M β-mercaptoethanol) and stored at −80°C at least overnight.

The RNA extraction protocol previously described was used (*76*). On the day of extraction, the frozen GTC-treated pellets were thawed at 37°C, harvested by centrifugation, resuspended in prewarmed triazole, and transferred to 2ml O-ring tubes containing sterile 0.2 mm zirconia beads. The triazole mixture was then bead-beaten twice with an intensity of 6 meters per second for 45 seconds, followed by the addition of chloroform and vigorous shaking for one minute. The samples were then centrifuged, and the aqueous layer removed and placed in Qiagen RNeasy columns for RNA cleanup. Genomic DNA was removed from all samples by performing a DNase digest on the columns before elution in pre-warmed RNase-free water. The extracted RNA was then removed from the BSL-3 facility. Concentration and quality of RNA (260nm/280nm absorbance) were assessed using a nanodrop spectrophotometer; samples were then stored at −80°C until submission. Samples were submitted to the Microbial ‘Omics Core (MOCP) and Genomics Platform at the Broad Institute of MIT and Harvard. QC, library preparation, ribodepletion, and paired-end sequencing on the Illumina NovaSeq 6000 following a modified RNAtag-Seq method (*77*) was performed by the MOCP.

### RNA-seq data processing and analysis

Sequencing reads were aligned to the *Mycobacterium tuberculosis* strain. Erdman = ATCC 35801 (RefSeq assembly accession: GCF_000350205.1) using Geneious Prime software (version 2023.2.1). The Geneious Prime software was used for the gene-level counts, and DESeq2 was used to identify genes that were differentially expressed in the acidic and neutral conditions. To identify acidic condition-responsive genes that were similarly differentially expressed across each strain, the up- and down-regulated gene lists were filtered by the criterion p < 0.05. Genes that were significantly differentially expressed and were regulated in the same direction (up or down) between all three strains were submitted for pathway and gene regulator enrichment analysis. To quantify enrichment p-values, a Fisher’s exact test was performed in R using fisher.test and a custom pathway and gene regulator annotation file that was curated through literature searches.

## Acknowledgments

We thank members of the Aldridge lab, Michelle Logsdon, and Robert Abramovitch for helpful discussions.

## Author contributions

Conceptualization: ESC, WCJ, MK, MM, ST, BBA

Methodology: ESC, WCJ, MK, TS

Experiment: ESC, WCJ, MK

Strain sharing: SV, NTTT

Writing – original draft: ESC, WCJ, MK, BBA

Writing – review & editing: all authors

