## Supplementary Materials for "Increased proportion of growth-arrested bacilli in acidic pH adaptation promotes *Mycobacterium tuberculosis* treatment survival"

### *Mycobacterium tuberculosis* adapts to acidic pH by increasing the proportion of growth-arrested bacilli, promoting survival to treatment

Eun Seon Chung *et al.*

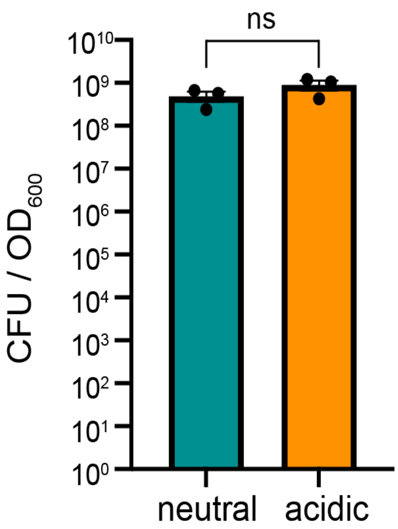


**Fig. S1. Cell viability quantification of Mtb adapted to neutral or acidic conditions.** Cell cultures were subjected to four days of pH adaptation, density quantified ($OD_{600}$) and plated for CFU (n = 3 biological replicates). Significance was determined by an unpaired Student's t-test: (p-value = 0.21).

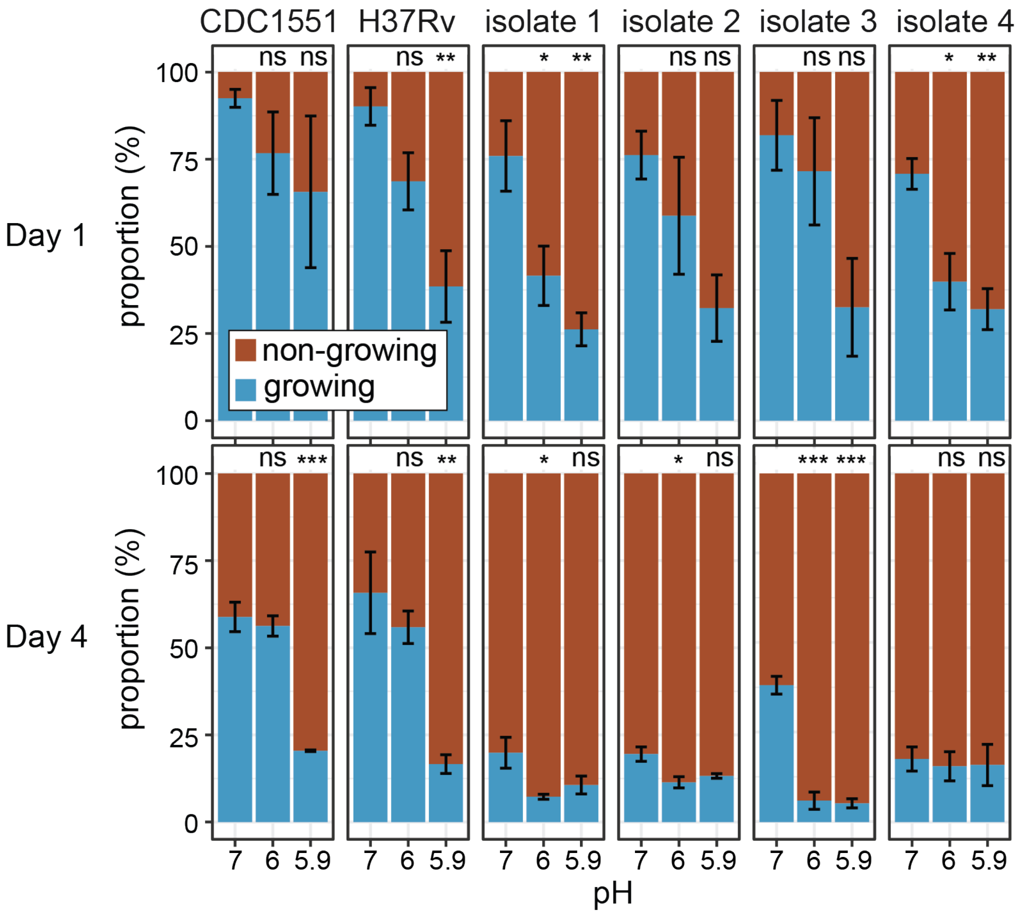


**Fig. S2. Proportion of growing and non-growing cells across six strains after pH adaptation.** The proportions of RADA-negative (non-growing) and RADA-positive (growing) cells are shown after one or four days of adaptation to neutral (pH 7.0) and acidic (pH 6.2 and pH 5.9) conditions. Among the six strains, two are Mtb laboratory strains (CDC1551 and H37Rv), and the other four are more recent clinical isolates (n = 3 biological replicates, error bars indicate the SEM). Significance was measured with one-way ANOVA and Dunnett's post-hoc test against neutral pH: ns, $p \geq 0.05$, *$p < 0.05$, **$p < 0.01$, ***$p < 0.001$.

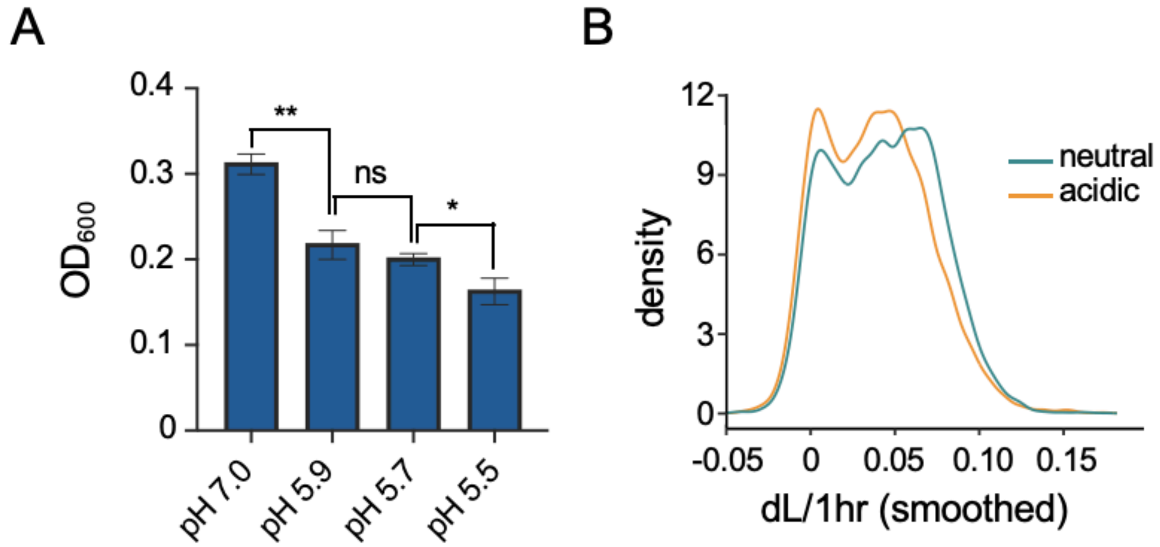


**Fig. S3. Slowed growth of Mtb at low pH is controlled by bimodal rate tuning.** (**A**) Density ($OD_{600}$) of Mtb cultures adapted to a range of neutral and acidic pH levels. p values, pH 7.0 vs. pH 5.9, p = 0.0014; pH 5.9 vs. pH 5.7, p = 0.18; pH 5.7 vs. pH 5.5, p = 0.019 (n = 3 biological replicates, error bars indicate the SEM). (**B**) Kernel density estimate of the growth rates (change in length per hour) of Mtb cells across 5 days of neutral (green) or acidic (orange) adaptation.

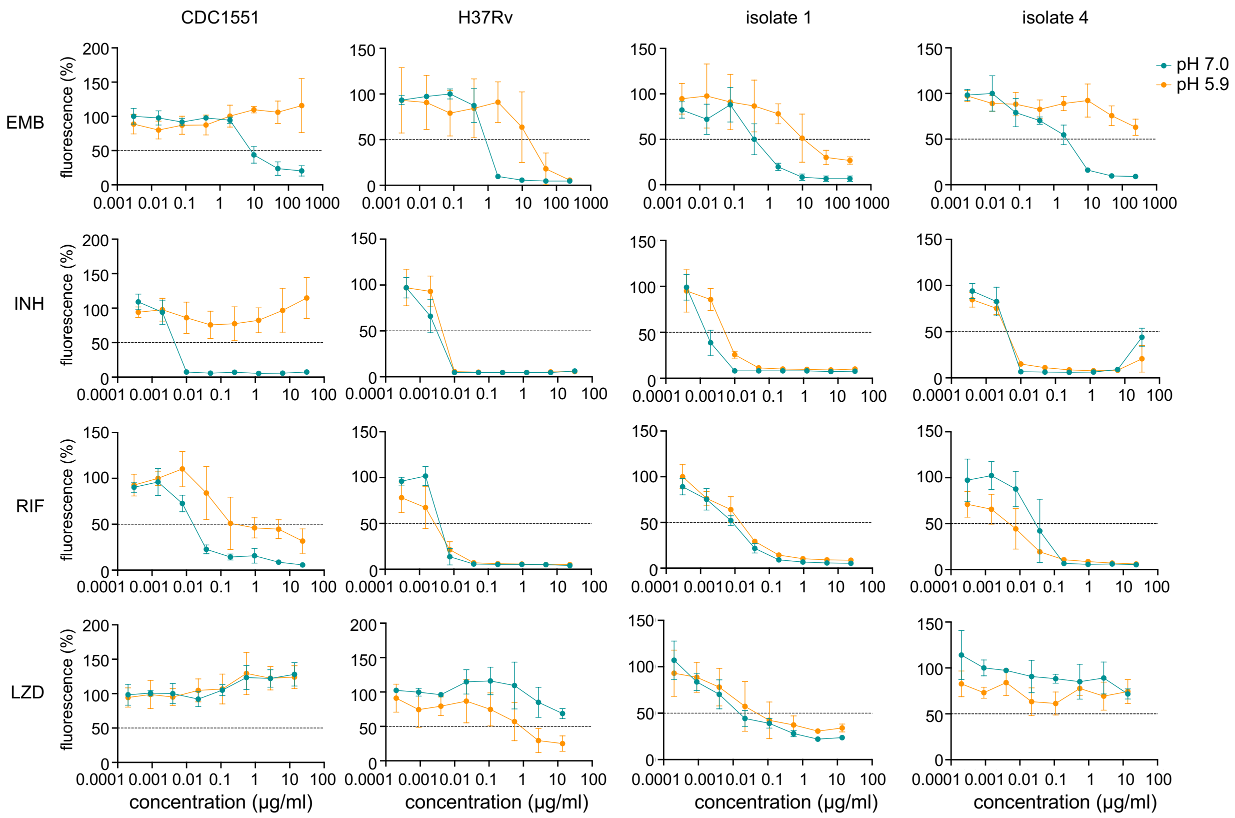


**Fig. S4. Survival of Mtb adapted to acidic or neutral conditions after 2-day treatment with TB antibiotics.** Four strains were used in the resazurin assay, including two laboratory strains and two clinical isolates. Ethambutol (EMB), isoniazid (INH), rifampicin (RIF), and linezolid (LZD) were added at various concentrations. Cells underwent ten days of drug treatment followed by ten days of recovery after drug removal. Following the recovery period, resazurin was added and incubated for one hour, after which fluorescence was measured to assess viability. Green indicates the neutral-adapted strain, and orange indicates the acidic-adapted strain. The resazurin fluorescence levels at the two pH values were normalized, with untreated strains set to 100%. Data are presented as the mean with SEM (n = 3 biological replicates).

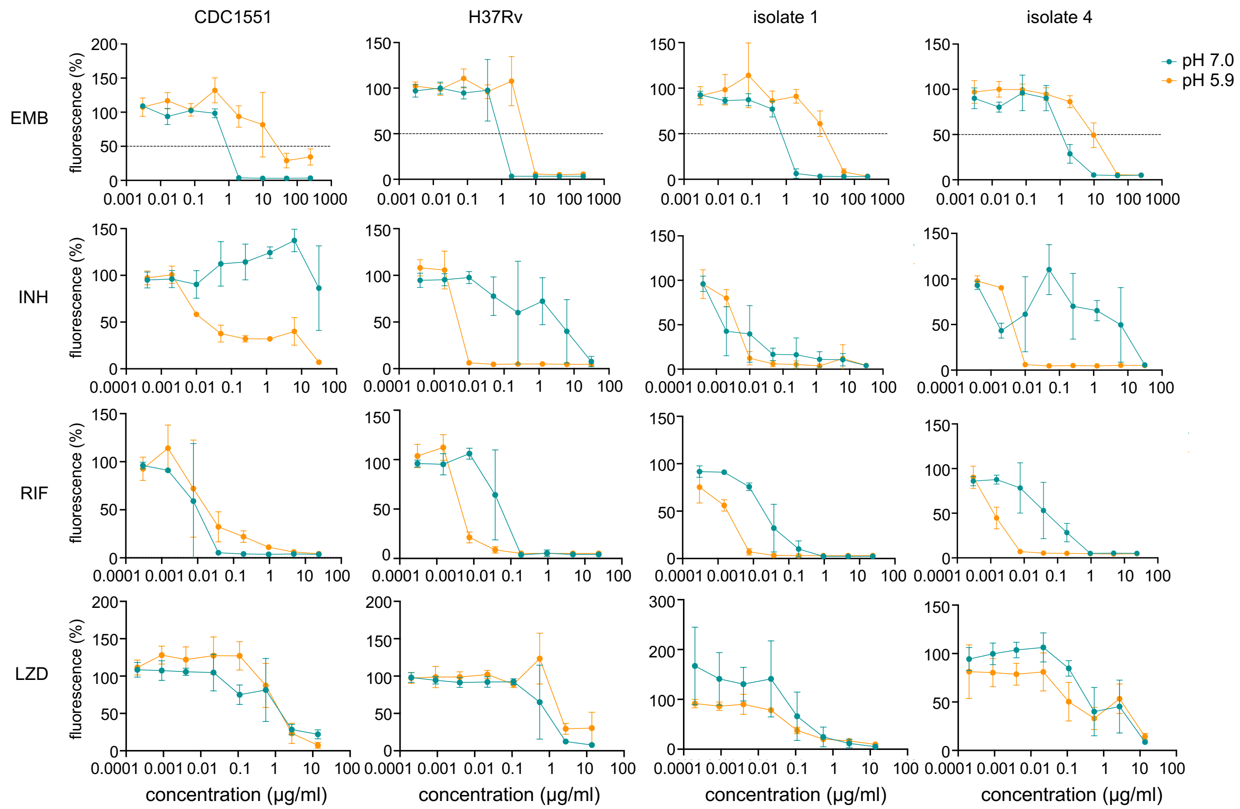


**Fig. S5. Survival of Mtb adapted to acidic or neutral conditions after 10-day treatment with TB antibiotics.** Four strains were used in the resazurin assay, including two laboratory strains and two clinical isolates. Ethambutol (EMB), isoniazid (INH), rifampicin (RIF) and linezolid (LZD) were added at various concentrations. Cells underwent ten days of drug treatment followed by ten days of recovery after drug removal. Following the recovery period, resazurin was added and incubated for one hour, after which fluorescence was measured to assess viability. Green indicates the neutral-adapted strain, and orange indicates the acidic-adapted strain. The resazurin fluorescence levels at the two pH values were normalized, with untreated strains set to 100%. Data are presented as the mean with SEM (n = 3 biological replicates).

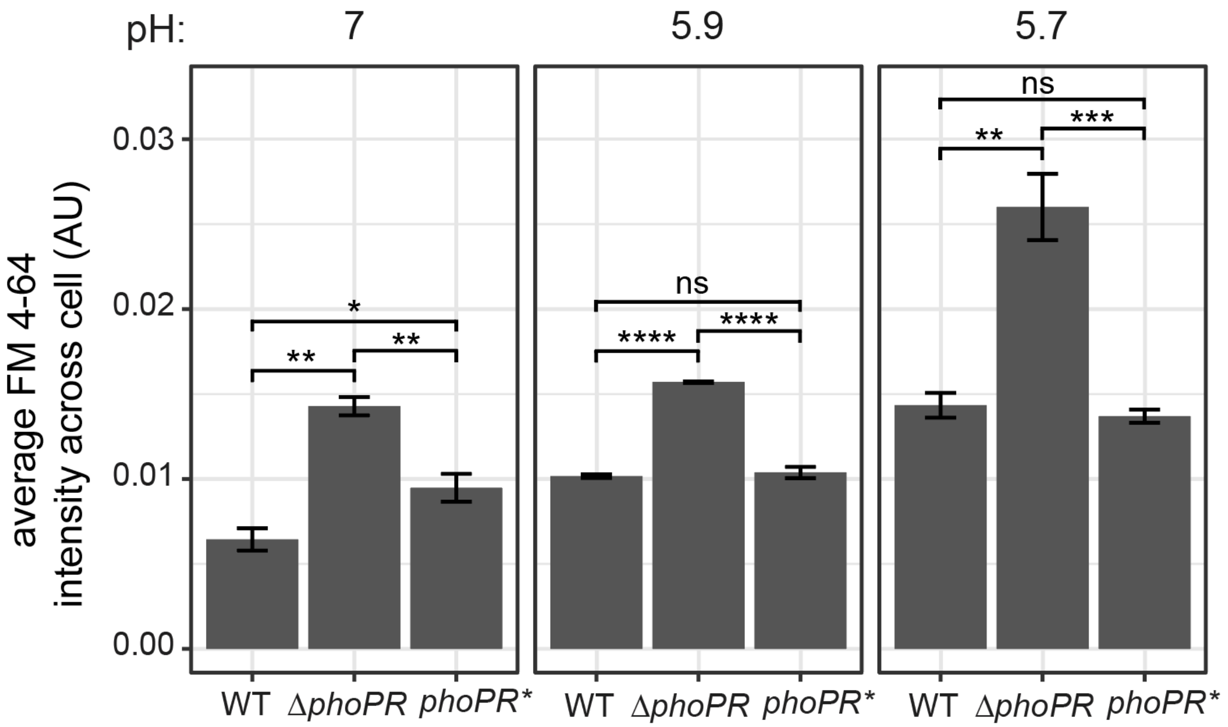


**Fig. S6. Deletion of PhoPR increases cell permeability at both neutral and acidic pH.** The CDC1551 WT, Δ*phoPR*, and *phoPR** strains were adapted to pH 7, pH 5.9, and pH 5.7 for one day and subsequently fixed and stained with FM 4-64 fluorescent dye. Mean and standard error are shown (n = 3 biological replicates). Significance was assessed with a one-way ANOVA and Tukey's post-hoc test: ns $p \geq 0.05$, *$p < 0.05$, **$p < 0.01$, ***$p < 0.001$, ****$p < 0.0001$.

| | Cells recovered | Cells not recovered | Total cells counted | Percentage of cells that recovered |
|---|---|---|---|---|
| HADA-positive | 5 | 731 | 736 | 0.7 |
| HADA-negative | 20 | 132 | 152 | 13.2 |
| total | 25 | 863 | 888 | |

**Table S1. Recovery quantification of EMB-treated Mtb.** Growth was determined based on HADA positivity (growing) or negativity (non-growing), and recovery was marked by an increase in cell length during the drug-free recovery period. Results are from one experiment (Fisher's exact test p-value = $3.8 \times 10^{-12}$).

| CDC1551 adjusted p-value | Isolate 1 adjusted p-value | Isolate 2 adjusted p-value | Gene Name | Rv# | log2 FC (fitness) |
|---|---|---|---|---|---|
| 2.8x10$^{-23}$ | 2.3x10$^{-26}$ | 3.4x10$^{-06}$ | Rv0412c | Rv0412c | -2.3 |
| 5.5x10$^{-69}$ | 3.1x10$^{-51}$ | 7.1x10$^{-22}$ | *pknG* | Rv0410c | -2.1 |
| 6.4x10$^{-12}$ | 7.4x10$^{-13}$ | 2.1x10$^{-07}$ | *embB* | Rv3795 | -2 |
| 1.1x10$^{-43}$ | 1.0x10$^{-21}$ | 4.0x10$^{-23}$ | Rv0996 | Rv0996 | -1.7 |
| 2.6x10$^{-10}$ | 2.1x10$^{-34}$ | 3.2x10$^{-11}$ | *leuC* | Rv2988c | -1.3 |
| 1.7x10$^{-111}$ | 5.0x10$^{-111}$ | 9.3x10$^{-24}$ | *mmpL8* | Rv3823c | -1.2 |
| 1.3x10$^{-12}$ | 3.1x10$^{-30}$ | 3.0x10$^{-12}$ | *nuoN* | Rv3158 | -1.2 |
| 1.4x10$^{-17}$ | 7.5x10$^{-127}$ | 5.5x10$^{-30}$ | *clpB* | Rv0384c | -1.1 |
| 6.8x10$^{-11}$ | 8.7x10$^{-38}$ | 3.1x10$^{-16}$ | *nuoM* | Rv3157 | -1 |
| 6.0x10$^{-17}$ | 1.2x10$^{-16}$ | 1.1x10$^{-16}$ | Rv1815 | Rv1815 | -1 |

**Table S2. pH-responsive genes that synergize with ethambutol when knocked down.** The intersection of the top 500 differentially expressed genes across all 3 strains were sorted by according to their fitness decrease (log2 fold change) upon CRISPRi knock down (*78*). The top 10 genes are displayed.
